# The evolution of sex is tempered by costly hybridization in *Boechera* (rock cress)

**DOI:** 10.1101/2020.02.11.944256

**Authors:** Catherine A. Rushworth, Tom Mitchell-Olds

## Abstract

Even after decades of research, the evolution of sex remains an enigma in evolutionary biology. Typically, research addresses the costs of sex and asexuality to characterize the circumstances in which one reproductive system is more favorable. Yet surprisingly few studies address the influence of common traits that are obligately correlated with asexuality, including hybridization and polyploidy; even though these traits have substantial impacts on selective patterns. In particular, hybridization is well-known to alter trait expression; these alterations may themselves represent a cost of sex. We examined the role of reproductive isolation in the formation of de novo hybrid lineages between two widespread species in the ecological model system *Boechera*. Of 664 crosses between *Boechera stricta* and *Boechera retrofracta*, 17% of crosses produced F1 fruits. This suggests that postmating prezygotic barriers, i.e. pollen-pistil interactions, form the major barrier to hybrid success in this system. These interactions are asymmetrical, with 110 F1 fruits produced when *B. stricta* was the maternal parent. This asymmetry was confirmed using a chloroplast phylogeny of wild-collected *B. stricta*, *B. retrofracta*, and hybrids, which showed that most hybrids have a *B. stricta* chloroplast haplotype. We next compared fitness of F2 hybrids and selfed parental *B. stricta* lines, finding that F2 fitness was reduced by substantial hybrid sterility. Our results suggest that multiple reproductively isolating barriers likely influence the formation and fitness of hybrid lineages in the wild, and that these costs of hybridization likely have profound impacts on the costs of sex in the natural environment.

## Introduction

Among the greatest outstanding puzzles in evolutionary biology is understanding the evolutionary processes that underlie the predominance of sex. A substantial swath of research focuses on population genetic models balancing the costs of sex and asexuality. Such costs are often framed as a comparison of sexual and asexual reproduction in constant environments, where sexual organisms produce two independent sexes. For this reason, these traditionally include the cost of producing independent male organisms (the “twofold cost of sex”), the costs of finding and attracting a mate, and the cost of recombination among co-adapted alleles (Maynard Smith, 1978; Lehtonen *et al.*, 2012). Simultaneously, costs of asexuality generally include clonal interference (Gerrish and Lenski, 1998) and Muller’s Ratchet (Müller, 1964), which rely on mutation accumulation without recombination.

The tension among the various costs of sexual and asexual reproduction is complicated by biological and ecological circumstances. Environmental variation has profound consequences for key elements of reproductive system, including fitness (Orr, 2009), gene flow (Richardson *et al.*, 2014), and recombination rate (Bomblies *et al.*, 2015; Modliszewski and Copenhaver, 2017). Previous studies show that ecological variation, in addition to population structure, contributes significantly to the maintenance of multiple reproductive modes (Agrawal, 2009; Becks and Agrawal, 2010, 2012). Yet despite the clear relevance of ecology for key elements of reproduction, few studies of the evolution of sex are conducted in the field (Neiman *et al.*, 2018). Additionally, numerous asexual systems are hybrids (e.g., Vrijenhoek, 1978; Lutes *et al.*, 2010; Coughlan *et al.*, 2017), suggesting that hybridization often causes, or at least co-occurs with, asexuality. A substantial body of literature examining organisms across the tree of life shows widespread effects of hybridization on fitness-related traits (Rieseberg *et al.*, 1999; Abbott *et al.*, 2013; Yakimowski and Rieseberg, 2014). The phenotypes altered by hybridization will have profound downstream effects on sexual/asexual dynamics. Yet disentangling the effects of asexuality from those of biological traits like hybridization is largely intractable, as these factors are tightly associated.

An additional complication stems from the intrinsic biological characteristics of sexual/asexual systems, and resulting variation in the applicability of the traditional costs of sex to all study systems. In flowering plants, asexual reproduction is widespread; apomixis (asexual reproduction via clonal seed) is found in nearly 20% of angiosperm families (Hojsgaard *et al.*, 2014). Most angiosperms produce flowers that contain both male and female organs, termed “hermaphroditic” (Barrett, 2002; Charlesworth, 2006). Thus, the framework for the costs of sex, namely the assumption of separate male and female organisms and a lack of reproductive assurance in sexuals, may be of limited relevance.

But additional costs of sex may exist, some of which offer promising avenues for research in flowering plants (Meirmans *et al.*, 2012). Chief among these is a fitness reduction associated with mating system, i.e. inbreeding and/or outcrossing (Charlesworth, 2006). Mating among related organisms, including self-fertilization, may reduce fitness (inbreeding depression, Charlesworth and Willis, 2009), while outcrossing between widely divergent individuals may likewise reduce fitness via outbreeding depression (e.g., Waser and Price, 1994). Although Meirmans and colleagues postulate that mating system negatively impacts sexuals, dissimilarity in the genesis of sexual and asexual lineages (i.e. hybrid origins of asexuals) suggest that the repercussions of mating system could also manifest in asexuals.

The influence of hybridization and mating system on the evolution of sex implicates the presence and expression of reproductively isolating barriers between species. While some barriers may prevent one species from encountering another (e.g., ecological/geographic prezygotic barriers), others hinder interspecific fertilization following mating (postmating prezygotic barriers). Postzygotic barriers render hybrids inviable or sterile via chromosomal rearrangements, cytonuclear interactions or Bateson-Dobzhansky-Muller incompatibilities (Coyne and Orr, 2004). Pollen-pistil interactions are a common form of postmating prezygotic barrier in flowering plants. These may be symmetrical, where each direction of a given cross is equally unable to achieve fertilization; or asymmetrical, where one direction of a cross succeeds, but the other fails (Tiffin *et al.*, 2001). Importantly, multiple barriers often co-occur, bolstering incomplete reproductive isolation caused by a single mechanism (Coyne and Orr, 2004).

The mustard genus *Boechera* is a widespread North American wildflower that engages in both sexual and asexual reproduction (Böcher, 1951; Al-Shehbaz and Windham, 1993). *Boechera* is highly self-fertilizing when sexually reproducing, which enables large-scale assessment of reproductive mode in interspecific hybrids via characteristic levels of heterozygosity (Beck *et al.*, 2012; Li *et al.*, 2017). Apomixis is widespread in *Boechera*, and co-occurs with hybridization or with outcrossing among divergent intraspecific populations (Rushworth *et al.*, 2018). The causative relationship between hybridization and asexuality is unknown, although apomixis may be a result of metabolic dysregulation caused by hybridization (Carman, 1997; Sharbel *et al.*, 2010). In the field, fitness of asexual lineages is higher than sexual lineages (Rushworth *et al.*, 2019), leading to two potential scenarios: first, uniformly high fitness of hybrid lineages that frequently reproduce asexually; or rampant but maladaptive hybridization events with rare transitions to asexuality “rescuing” hybrid genotypes. Previous studies of ecological and genetic variation in *Boechera stricta* suggest that reproductive isolation between subgroups is driven by ecological differentiation (Lee and Mitchell-Olds, 2011, 2013) and a single chromosomal inversion underlies phenological divergence (Lee *et al.*, 2017). But to date, little is known of postmating barriers in this group.

Here we explore three vignettes investigating the costs of hybridization in the evolution of sex in *Boechera*. We found that the formation of hybrids between the two best-characterized species, *Boechera stricta* and *Boechera retrofracta*, is hindered by the expression of multiple reproductive isolating barriers. The main barrier to hybrid success occurred before hybrids were formed, with most F1 crosses failing to set seed. This barrier was asymmetrical, with substantially higher success for F1s with *B. stricta* as the maternal parent than the reverse cross. Phylogenetic analysis of a chloroplast marker confirmed that *B. stricta* most often acts as the maternal parent in wild-collected hybrids. We next compared fitness of hybrid F2s with their selfed parental *B. stricta* lines, finding that hybrid sterility substantially reduces F2 fitness, although fertile F2s produce more fruits than selfed lines. These results have important implications for the speciation process and for the evolution of sex in this ecological model system.

## Materials and Methods

### Crosses and plant growth

In the wild, *B. stricta* and *B. retrofracta* commonly co-occur, hybridize, and form asexual lineages (Rushworth *et al.*, 2018). In 2012, one line was selected from 11 populations of each species for use as crossing parents (Figure 1, Table S1 in Supplementary Material). In five populations, both species co-occurred in close proximity to one another; these populations are thus considered sympatric. All lines were of known genotype from previous studies (Song *et al.*, 2006; Lee and Mitchell-Olds, 2011; Rushworth *et al.*, 2018). Each line had been previously grown in the greenhouse for 1-2 generations. Seeds were germinated on wet filter paper in petri dishes and grown in greenhouses at Duke University, then vernalized for 6 weeks at 4°C with 12 hour daylength. We used three staggered planting cohorts to allow for genotypic variation in phenology.

**Figure 1.**
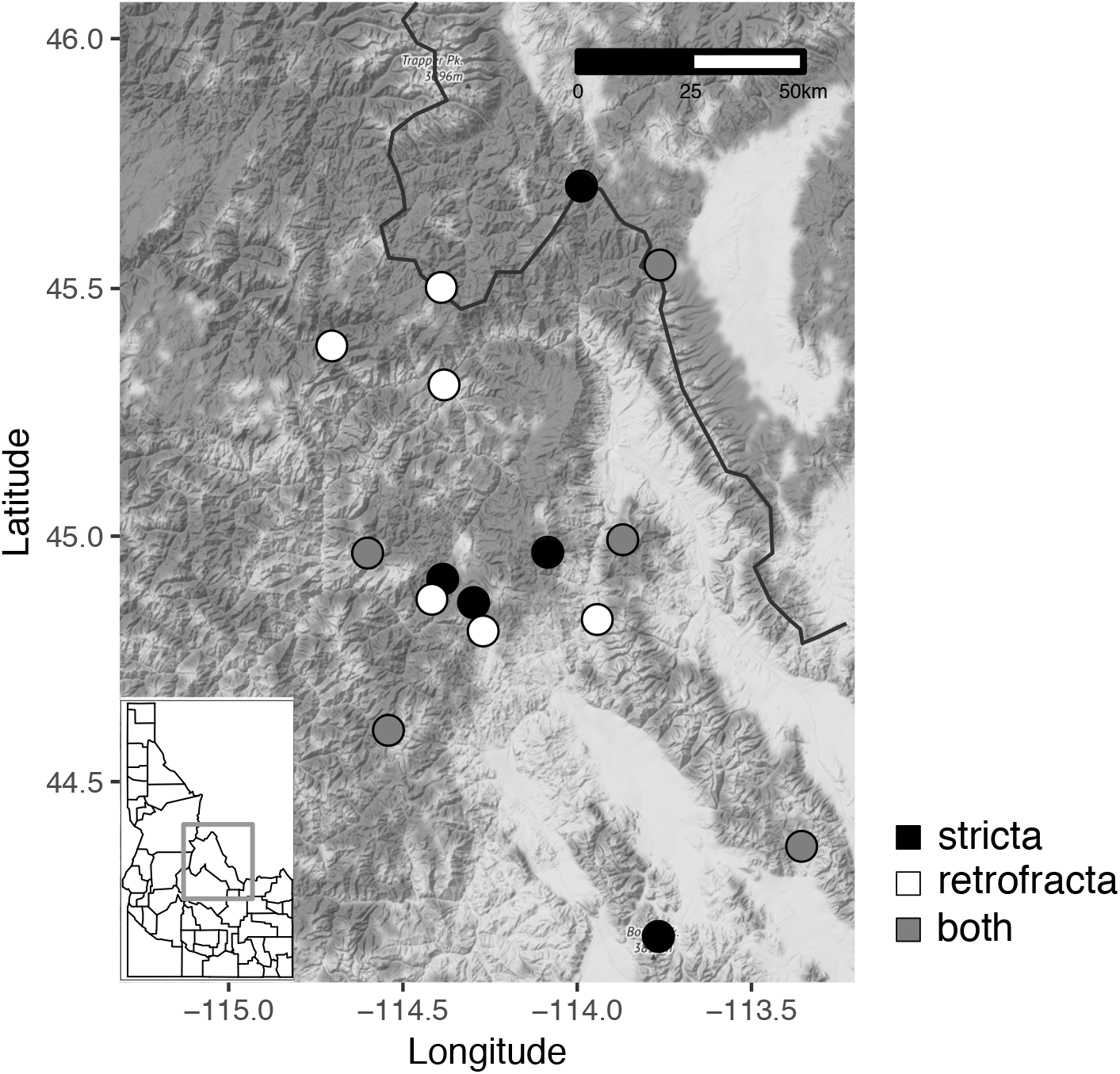
Map of study populations used as cross parents. Each circle represents a population. *B. stricta* populations are represented by filled circles, and *B. retrofracta* populations are represented by unfilled circles. Populations where *B. stricta* and *B. retrofracta* co-occur are indicated by gray fill. Plants were collected in central Idaho and western Montana (inset). One population, HES, was collected in Colorado and is not shown.

Plants were subsequently moved to growth chambers in Duke University’s Phytotron. Growth conditions were 22°C constant temperature with ambient relative humidity and carbon dioxide, and 12 hour days at 350 micromoles of light. Reciprocal crosses were performed on two to five flowers for each possible parental combination following Schranz et al. (2005). Upon observation of cross failure, replicate crosses using individuals from different cohorts were made. Concurrently, other flowers on the same plants were permitted to self-fertilize. Additional flowers were self-fertilized by hand.

Fruiting success was calculated as the number of fruits divided by the number of flowers for a given cross type (maternal *B. stricta*; paternal *B. stricta*; hand-selfed *B. stricta*; hand-selfed *B. retrofracta*). To correct for differences in fruit set inherent to each species, we also calculated crossing indices for each cross type (McDade and Lundberg, 1982). Crossing indices were calculated by dividing fruiting success of an interspecific cross by fruiting success of the hand-selfed maternal parent (i.e., maternal *B. stricta* crosses divided by hand-selfed *B. stricta*; paternal *B. stricta* crosses divided by hand-selfed *B. retrofracta*).

When F1s produced <15 seeds, we germinated all available F1 seeds; if a given parental combination resulted in >15 seeds, we germinated 15 seeds. All F1 seeds were germinated on filter paper in petri dishes and grown in greenhouses at Duke University. All seeds available, regardless of germination status, were planted onto a mix of MetroMix 200 and Fafard 4P Mix (Sun Gro Horticulture, Agawam, MA, USA) in Ray Leach Conetainers (Steuwe and Sons, Tangent, OR, USA).

To verify that F1s were the product of successful crosses, viable F1s were genotyped at three variable microsatellite loci (*ICE3*, *c8*, *BF20*; Clauss *et al.*, 2002; Dobeš *et al.*, 2004a; Song and Mitchell-Olds, 2007) following protocols from Beck et al. (2012). This resulted in 50 viable unique F1 seed lines. Selfed parental lines were also genotyped to verify self-fertilization. To verify reproductive mode of F1s, 3—12 (mean 9.1) F2 individuals were genotyped in the same manner as F1s. A given plant was determined to be sterile if it failed to reproduce in three months, one month longer than the typical growing season in the central Rocky Mountains.

Poor F2 seed set resulted in a final experimental total of seven unique F2 families, derived from the same *B. retrofracta* parent and four *B. stricta* individuals (Table S1). Parental lines were also permitted to self for a second generation. Segregation of alleles in all lines was observed, indicating that F1s reproduced sexually.

### Greenhouse experiment

Fitness of F2s and their selfed parental lines was assessed in the greenhouse in 2014. Seeds were germinated on wet filter paper in petri dishes and transplanted as seedlings. Low germination for some F2 families resulted in an unbalanced design. In total, 573 individuals (300 parents, 273 F2s) were used, with 60 replicates from each F1 parental line and 19-59 progeny for each F2 family (mean 39 replicates). 52 individuals (51 hybrid F2s and one selfed parental individual) died during the course of the experiment.

Plants were grown in randomized blocks and racks were rotated to minimize microhabitat variation. Traits measured included those related to fitness (probability of survival and reproduction, flower number, fruit number, aborted fruit number) and biomass (rosette width, plant height, leaf number). Aborted fruits were those that contained only partially developed (i.e. inviable) seeds. Average seed set was calculated from four replicates of each genotype; for genotypes EJ1 and EJ2, less than four individuals reproduced, resulting in seed set estimation from one and two individuals, respectively. Total fitness was calculated as the number of fruits per individual multiplied by average seed set per genotype, with all zeros included. Fecundity was calculated as the number of fruits per individual multiplied by average seed set per genotype, conditional on reproduction.

### Statistical analyses

To understand how cross type influenced multiple fitness components, we used generalized linear mixed models (GLMMs). We estimated each fitness component (probability of reproduction, fruit number, aborted fruit number, seed set, and total fitness) as a function of cross type (hybrid or selfed), with a scaled covariate of rosette width in each model to account for the impact of plant size on reproductive output. Because all F2s had the same paternal *B. retrofracta* genotype, this genotype was constant; we thus compared four *B. stricta* lines with their hybrids. All models included random effects for experimental block and parent nested in cross type. In four of five models, a random effect of *B. stricta* parent line was also included; the model for probability of reproduction estimated the variance among parents to be zero, and thus this term was eliminated from the model. An additional random effect of genotype (F2 family or parental genotype) was incorporated in the total fitness model, as it had a substantial effect on model fit.

Models for fruit number, aborted fruit number, and fecundity used a negative binomial error distribution, while a binomial error distribution was used for the probability of reproduction. A single outlier that produced 308 fruits was removed from the fruit number, aborted fruit number, and total fitness models. Lifetime fitness was estimated with a zero-inflated negative binomial distribution using the canonical link functions. In most fitness models, structural zeros account for plants that failed to survive. Because survival is not a factor in controlled conditions, structural zeros in the model indicate plants that failed to reproduce, while the conditional portion of the model represents the fecundity of the plants that did reproduce. Zero-inflation was modeled across both main effects.

All analyses were run in R version 3.5.2. Directionality of cross success via F1 fruit production was analyzed using Fisher’s exact tests in the base R stats package. Due to very small sample sizes, statistical comparison of success was not possible for later stages of hybrid development. GLMMs were run using the packages lme4 (probability of reproduction model; Bates *et al.*, 2015) and glmmTMB (all other models; Brooks *et al.*, 2017). Significance was calculated via likelihood ratio tests. *P*-values were adjusted using the Bonferroni-Holm method. Estimated marginal means were calculated with the package ggeffects (Lüdecke, 2018).

### Molecular phylogenetics

We used 112 samples of *B. stricta*, *B. retrofracta*, and *B. retrofracta* × *B. stricta* in a molecular phylogenetic analysis. Samples from *B. retrofracta* (N=32) and *B. retrofracta* × *B. stricta* (N=31) were selected from across central Idaho and western Montana, while *B. stricta* samples (N=49) represent the full species range (Table S2). DNA was extracted from each sample using either Qiagen DNeasy Plant Mini Kits (Qiagen, Hilden, Germany), a modified CTAB protocol (Beck *et al.*, 2012), or following Lee et al. (2011).

To identify wild-collected hybrid parentage, we amplified the intron and second exon of the chloroplast gene *trn*L, using primers c and d and thermal cycling protocols from Dobes et al. (2004b). PCR was performed with 20 mL reactions consisting of 10uM of each primer, 0.2 uL of 10 mg/mL BSA (Millipore/Sigma), and 2mM dNTPs, 10X buffer and Choice-Taq DNA polymerase (all Denville Scientific, Metuchen, NJ, USA). PCR was run on MJ Research PTC-200 thermal cyclers and Sanger sequencing was performed at the UC Berkeley DNA Sequencing Facility on an ABI 3730xl analyzer (Applied Biosystems). *B. stricta* sequences were provided by Baosheng Wang via sequencing data in Wang et al. (2019).

A maximum parsimony phylogeny was inferred using Paup version 4.0a166 (Swofford, 2003). We performed a search from ten different random addition sequence starting trees, using TBR branch swapping and a reconnection limit of 8.

## Results

### Hybrid crosses rarely succeed

The success of hybrid crosses was limited at multiple developmental stages. We conducted 203 reciprocal crosses using unique parental combinations, with an average of 3.07 flowers (±0.04 standard error, se) for each cross, resulting in 664 total crossed flowers. Crosses had a 17.3% chance of successfully producing fruits. This resulted in 115 total F1 fruits from all maternal plants, an average of 0.53 fruits per flower (±0.06 se). Although *Boechera* fruits from autonomous self-fertilization may produce upwards of 100 seeds (Rushworth *et al.*, 2011), these F1 fruits produced an average of 12.98 seeds per fruit (±2.32 se), totaling 2804 F1 seeds from all crosses. Despite low average seed set, 59 of 115 F1 fruits produced >15 F2 seeds.

To assess seed viability, we planted 1190 F1 seeds, resulting in 829 germinants from 98 F1 fruits (an average of 1.25 seeds ±0.14 se per cross). Seventeen fruits produced no viable seed. This represented a 70% rate of viability. 705 seedlings from these 98 F1 lines were planted onto soil, but in 77 lines, all germinants perished. 88 plants from 21 F2 lines survived. Of these 88 plants, 59 plants from 13 lines reproduced. The remaining plants were sterile. Genotyping of F2 seeds showed that six of these 13 lines were not successful crosses (i.e., they were identical to the maternal genotype), leaving seven total F2 crossing families for the greenhouse experiment.

### Directionality of cross affects success

*B. stricta* was the maternal parent for all seven successful F2 lines used in our experiment. The influence of cross directionality on success became apparent early on. Despite conducting a roughly equivalent number of crosses in each direction (317 flowers with *B. stricta* as maternal parent, 347 paternal *B. stricta* crosses), only five of 115 F1 fruits, representing replicates of 62 unique parental combinations, were produced by crosses with *B. stricta* as the paternal parent. We next compared the proportion of unique parental combinations that resulted in fruits with a Fisher’s exact test. 57 of 102 maternal *B. stricta* parental combinations resulted in fruits, compared to 5 of 101 paternal *B. stricta* combinations. These proportions are significantly different, with maternal *B. stricta* crosses 23.9 times as likely to result in fruits as paternal *B. stricta* crosses (Fisher’s exact test, 95% CI 8.84—81.53, *P*=2.199e-16; Figure 2).

**Figure 2.**
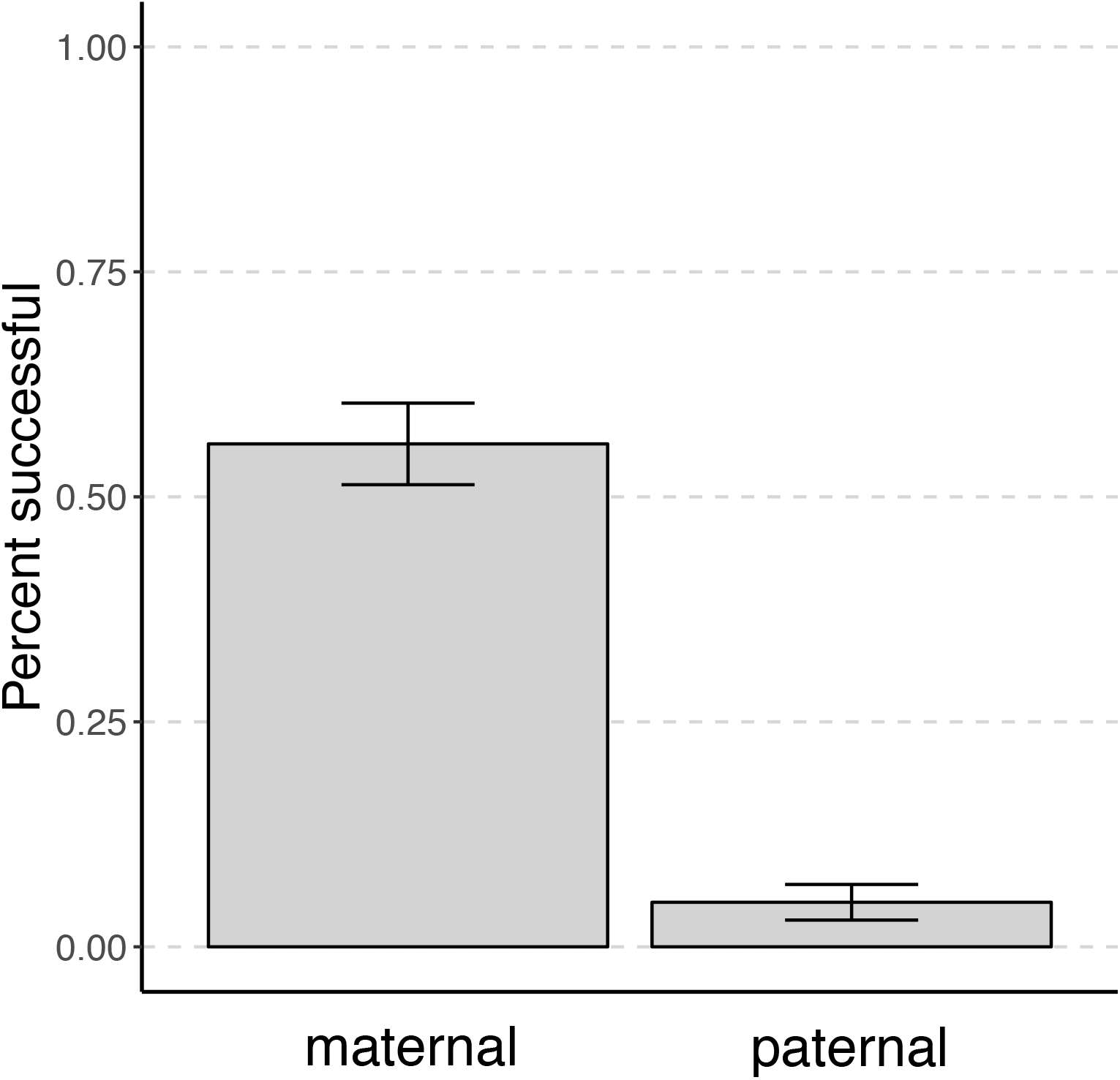
Pollen-pistil barriers are asymmetrical. Interspecific crosses with *B. stricta* as the maternal parent were more likely to set seed than those with *B. stricta* as the paternal parent. Proportion of unique parental combinations that resulted in F1 fruit are shown. Error bars indicate 95% CIs.

We note that fruiting success was very low for hand-selfed *B. stricta* (8 fruits out of 37 crossed flowers, or 0.216 fruits/cross) and *B. retrofracta* (3 fruits out of 44 crossed flowers or 0.068 fruits/cross), suggesting mechanistic issues with crossing. By comparison, paternal *B. stricta* crosses produced 5 fruits out of 347 crossed flowers (0.014 fruits/cross) and maternal *B. stricta* crosses produced 110 fruits from 317 crossed flowers (0.347 fruits/cross). The crossing index for maternal *B. stricta* was thus 1.605 (0.347/0.216), while the crossing index for paternal *B. stricta* was 0.211 (0.014/0.068), consistent with higher fruiting success in maternal *B. stricta* crosses despite poor responses to hand-fertilization in both species.

Of five potential paternal *B. stricta* F1 lines, all reproduced. Four produced only one F2 seed, although one of these failed to germinate, while one produced 13 F2 seeds. Genotyping confirmed that two of these crosses, including the high seed set genotype, were not successful (i.e. the putative F1 was identical to the maternal parent). Of the two remaining crosses, one germinant died early, and the last resulted in a sterile plant. Thus, no paternal *B. stricta* crosses were ultimately successful in reproducing, suggesting *B. stricta* is only suitable as a maternal parent.

Because hybridization dynamics may strongly differ in the field, we used a phylogenetic analysis of the maternally-inherited chloroplast to assess cross directionality of wild-collected *B. stricta* × *B. retrofracta* hybrids. Our plastid *trn*L alignment of 112 accessions (49 *B. stricta*, 32 *B. retrofracta*, 31 hybrids) had 10 variable characters, five of which were parsimony-informative. The most parsimonious trees had 11 changes, and resolves two major groups, one containing 48 of 49 *B. stricta* accessions, and the other containing 28 of 32 *B. retrofracta* (Figure 3). 30 of the 31 hybrid accessions fell in the *B. stricta* group, with 28 of them sharing a haplotype with a sampled *B. stricta* accession. Although previous studies show chloroplast haplotypes are shared between the two species (Sharbel and Mitchell-Olds, 2001; Dobeš *et al.*, 2004b), our results indicate that *B. stricta* is usually the maternal parent in the wild.

**Figure 3.**
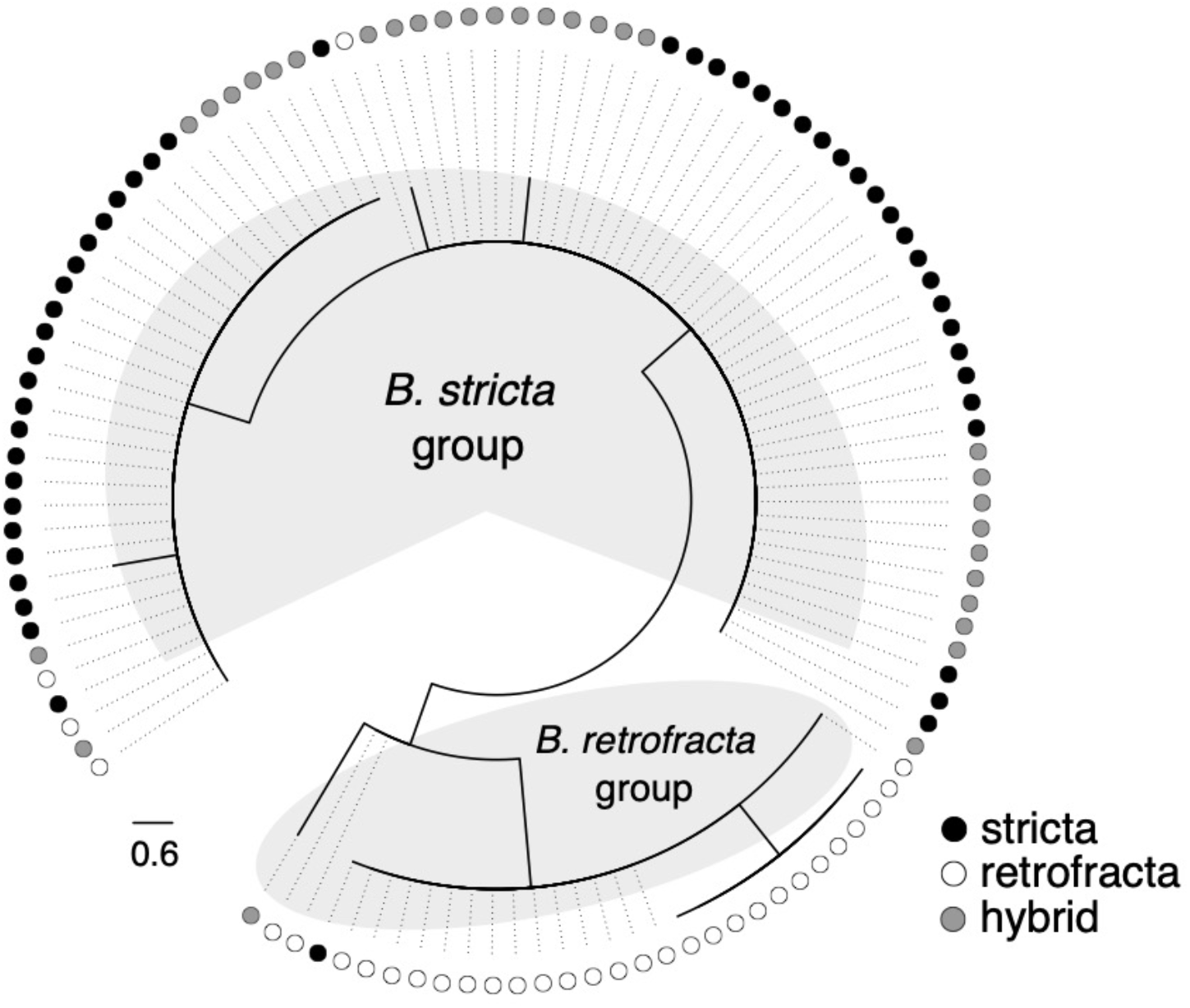
Phylogeny of chloroplast marker *trn*L. Sequenced taxa show a clear split between two groups, one of which is predominantly *B. stricta* (filled circle), and the other predominantly *B. retrofracta* (empty circle). 30 of 31 wild-collected *B. stricta* × *B. retrofracta* hybrids group with *B. stricta*, consistent with *B. stricta* acting as maternal parent in the majority of naturally-formed hybrids.

### Hybrid fitness is lower than selfed lineages

Total fitness (total maximum seed set, a product of the probability of reproducing and fecundity) was lower in F2 hybrids than in selfed lineages (overall significance, Table 1). On average, hybrids produced 510 (±43.3 se) seeds per plant, while selfed genotypes produced 784 (±30.6 se) seeds per plant. 67.8% of hybrids (±0.03 se) reproduced vs. 96.2% of selfed lines (±0.01 se). Hybrids were far less likely to reproduce than selfed lines, which drove this difference in fitness (zero-inflated model, Table 1, Figure 4A), although this difference was not significant following correction for multiple comparisons in the reproduction-only model (Table S3).

**Table 1.** Hybrids have lower total fitness due to reduced probability of reproduction. Results for the fixed effects of cross type (selfed vs. hybrid) and plant size from a zero-inflated negative binomial GLMM. Left, the zero-inflation portion models structural zeros, or plants that did not reproduce. Fewer hybrids reproduced than selfed parents. Center, the conditional portion of the model shows that cross type has no effect on seed set. Right, significance estimates for the overall model, incorporating both portions. Estimates (coef) and standard errors (se) come from conditional models, while test statistics (*χ*^2^ deviance, degrees of freedom, and *P* values) come from likelihood ratio tests for each overall effect. Results from independent models of reproduction probability, fecundity, and aborted fruit number are in Supplementary Information. Significant *P*-values are shown in bold.

|  |  |  | *Zero-inflation* |  |  |  | *Conditional* |  |  |  | *Overall* |  |  |
| --- | --- | --- | --- | --- | --- | --- | --- | --- | --- | --- | --- | --- | --- |
| term | condition | df | coef | se | $\chi^2$ | *P* | coef | se | $\chi^2$ | *P* | df | $\chi^2$ | *P* |
| Cross type | selfed | 1 | -2.74 | 0.419 | 9.02 | **0.019** | 6.63 | 0.252 | 0.32 | 0.65 | 2 | 9.34 | **0.047** |
|  | hybrid |  | -1.55 | 0.205 |  |  | 6.48 | 0.177 |  |  |  |  |  |
| Size | - | 1 | -2.65 | 0.182 | 46.91 | **4.47e-11** | 6.57 | 0.053 | 2.45 | 0.24 | 2 | 49.35 | **9.60e-11** |

**Figure 4.**
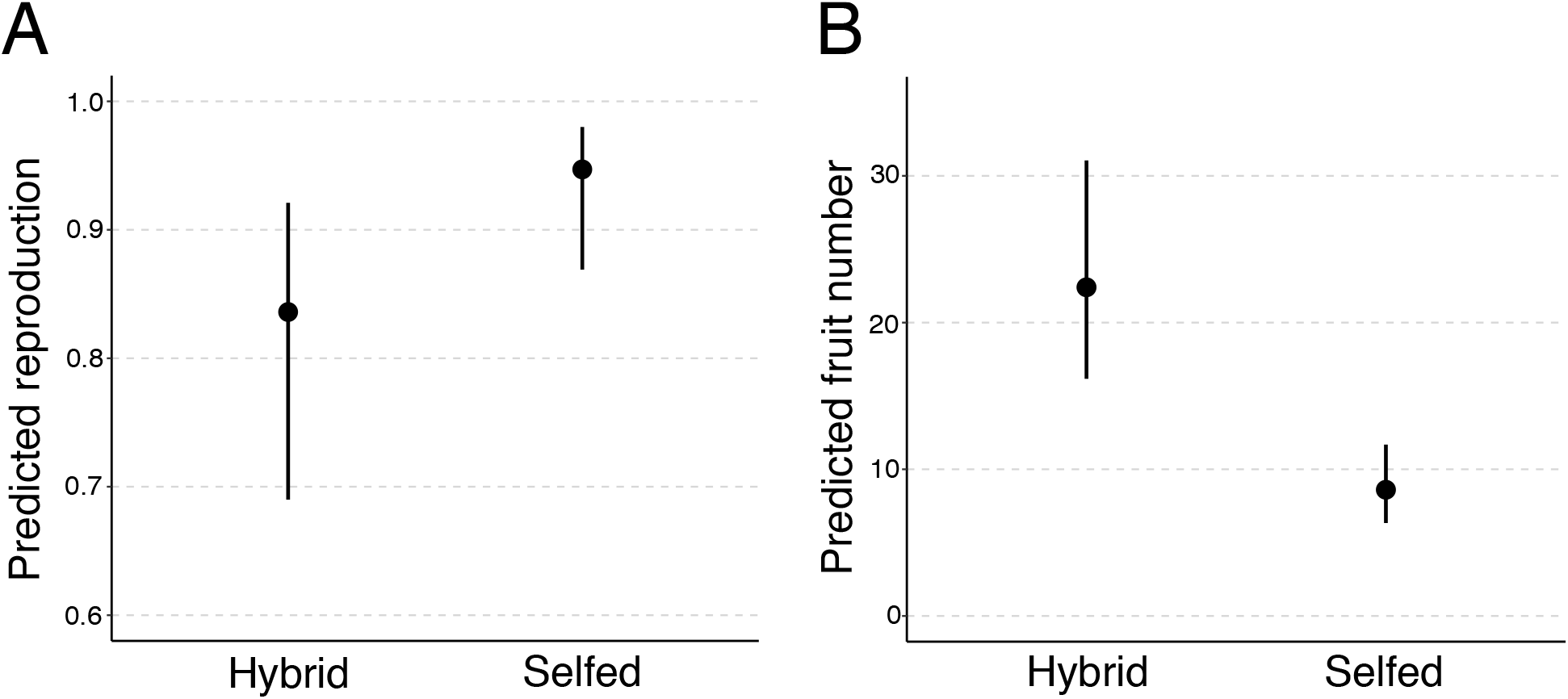
Hybrids are less likely to reproduce, but produce more fruits than sexual lineages. Estimated marginal means from GLMMs show the main effect of cross type on A) the probability of reproducing; B) fruit number. Bars show 95% confidence intervals.

However, F2 hybrids produced more fruits than selfed lines, averaging 22.5 ±1.62 se compared to 10.6 ±0.4 se for selfed lines (Figure 4B, Table 2). Hybrids had notably higher variance in fruit number, with a standard deviation of 24.1 fruits compared to 6.17 for selfed lines. Indeed, one hybrid individual produced 308 fruits, while the maximum number of fruits produced by a selfed line was 48. F2 hybrids also produced more aborted fruits than selfed lineages; this difference was significant prior to correction for multiple comparisons (*P*=0.02), but not after (Table S4). Fecundity, or seed set conditional upon reproduction, did not differ between hybrids and selfed lines, with hybrids producing an average of 709 seeds ±52.3 se compared to 814 ±30.0 se for selfed lines (conditional model, Table 1, Figure S1, Table S5). Estimated marginal means for all individual models are reported in Table S6.

**Table 2.** Hybrids produce more fruits than sexuals. Results for the fixed effects of cross type (selfed vs. hybrid) and plant size from a negative binomial GLMM. Estimates (coef) and standard errors (se) come from conditional models, while test statistics (*χ*^2^ deviance, degrees of freedom df, and *P* values) come from likelihood ratio tests for each overall effect. Significant *P*-values are shown in bold.

| term | condition | df | coef | se | $\chi^2$ | *P* |
| --- | --- | --- | --- | --- | --- | --- |
| Cross type | selfed | 1 | 2.09 | 0.221 | 8.98 | **0.019** |
|  | hybrid |  | 3.05 | 0.164 |  |  |
| Size | - | 1 | 3.48 | 0.077 | 28.86 | **3.12e-07** |

Selfed lines were not significantly larger than hybrid lines (hybrid mean 90.2mm ±1.95se vs. selfed mean 127mm ±1.02 se), although hybrid size was more variable (hybrid standard deviation, sd=29 vs. selfed sd=15.8; Figure S2). Size had a significant effect on total fitness (overall model, Table 1), probability of reproduction (zero-inflated portion, Table 1) and fruit number (Table 2). Notably, a random effect of genotype accounted for 14% of the variance in the total fitness model, far more than any other random term in this model or in any other (Table S7). This suggests that genotype plays a large role in the ultimate trajectory of hybrid lineages.

## Discussion

Hybridization is known to co-occur with transitions to asexuality in numerous plant systems (summarized in Asker and Jerling, 1992). Although a substantial body of research explores the evolution of sex via a balance between its costs and those of asexuality, few studies account for the effects of correlated traits. The strong link between asexuality and hybridization across the tree of life is likely to influence the evolution of sex in several unique ways. First, hybridization that occurs among sexual lineages and results in outbreeding depression may represent a cost of sexual reproduction. Alternatively, if hybridization is frequently associated with transitions to asexuality, but most asexual hybrids exhibit lower fitness, hybridization may be considered a cost of asexuality. Second, asexuality may act as a “rescue” of low fitness hybrids, enabling reproductive assurance for lineages that might otherwise go extinct. Third, hybridization alters phenotype expression in myriad ways, resulting in traits that are transgressive, intermediate, or novel. These alterations, in turn, will have strong ecological impacts on the evolutionary trajectories of hybrid lineages.

Here we explored the fitness consequences of de novo hybrids in controlled conditions. We found that the costs of hybrid formation may far outweigh any benefits that successful hybrid asexuals experience in the natural environment. F2 hybrids are less likely to reproduce, which results in reduced fitness (Table 1, Figure 4), despite producing more fruits than their selfed counterparts (Table 2). Hybrids are also difficult to form; hybridization events led to F1 fruit formation in less than 20% of crosses. Additionally, cross directionality is of profound importance. In our experiment, only hybrids with *B. stricta* as the maternal parent were ultimately successful (Figure 2). A chloroplast phylogeny corroborates this result, implicating asymmetrical reproductively isolating barriers in the formation of hybrids (Figure 3). Collectively, multiple reproductively isolating barriers reduce hybrid fitness, which has important consequences for hybrid lineages in the wild.

Numerous traditional costs of sex, such as the twofold cost of males and the metabolic costs of attracting a mate, have been discussed in the literature (Lehtonen *et al.*, 2012). Recently, Meirmans et al. (2012) proposed hybridization, inbreeding, and outcrossing, which we collectively will refer to as mating system, as an additional cost of sex. As conceived by the authors, outcrossing and hybridization will negatively impact the fitness of sexual lineages, via reproductive isolation or outbreeding depression. Similarly, inbreeding within small populations may reduce sexual fitness through inbreeding depression. Through this lens, the fitness consequences of mating system fall on sexual populations, and asexuals, once formed, are able to avoid fitness reductions by avoiding mating altogether.

However, mating system may instead pose a cost to asexuals. Additionally, whether mating system costs impact sexuals or asexuals may depend on other factors. For example, the nature of hybridization costs will depend on the frequency of transition to asexual reproduction following hybridization. If all hybrids transition to asexuality, the developmental issues caused by reproductively isolating barriers may hinder asexual formation and success. If the majority of hybrids are sexual, with rare transitions to asexuality, the fitness detriments of hybridization will pose a cost to sexual populations.

Knowledge of a system’s mating system is critical to attributing this cost. For example, frequent self-fertilization among sexual lineages will largely enable avoidance of the fitness costs of hybridization. While asexual *Boechera* result from outcrossing events between either populations or species, sexual lineages are highly self-fertilizing (Beck *et al.*, 2012; Li *et al.*, 2017; Rushworth *et al.*, 2018). Nevertheless, hybridization is clearly common. Large-scale collections-based efforts have identified 400 unique asexual hybrid combinations of *Boechera* in North American herbaria; the vast majority of these are asexual (Li *et al.*, 2017). This observation suggests two possible scenarios: either rare hybridization events always co-occur with asexuality; or rampant hybridization produces numerous short-lived sexual lineages, some of which transition to asexuality. With more knowledge of both the mechanisms of asexual formation, and the outcrossing and hybridization rate in natural populations, distinguishing between these possibilities offers a promising area for future research.

Hybridization may represent a cost of both sex and asexuality, through its substantial and varying influence on the expression of numerous phenotypic traits. Importantly, the mode of expression will depend on the genetic basis of the trait under consideration. Hybrids may produce traits that are intermediate to their parents, or the phenotype of one parent, or even transgressive or novel (Rieseberg *et al.*, 1999; Abbott *et al.*, 2013; Yakimowski and Rieseberg, 2014). Similarly, hybridization is likely to influence different components of fitness. Fitness is an inherently complex trait, putatively underlain by thousands of loci across the genome. Indeed, heterosis in maize is well-known to increase fitness, likely through the masking of many deleterious recessive alleles (Springer and Stupar, 2007). Simultaneously, certain fitness components may be simple. For example, overdominance at a single locus *SFT* in tomato causes production of indeterminate infructescences, vastly increasing the number of fruits (Krieger *et al.*, 2010). Importantly, the genotype of each parent will have a substantial impact on the nature of hybrid trait expression, as seen in our total fitness model. Further disentangling of the unique fitness impacts of hybridity and its ramifications for the evolution of sex is an open prospect in theory and experimental research, and *Boechera* is well-situated for this work.

Impediments to *Boechera* hybrid formation at multiple stages of development suggest the action of multiple reproductive barriers. Hybrid inviability (indicated by relatively low germination of hybrids) and sterility (indicated by both a failure to reproduce and by the production of more aborted, or sterile, fruits) are both factors in F2s, suggesting the presence of Bateson-Dobzhansky-Muller incompatibilities. However, the main barrier to hybrid formation in our crosses occurred prior to fertilization. The failure of most F1 crosses to set seed is consistent with postmating prezygotic barriers. Pollen-pistil barriers, when incompatible interactions between pollen and pistil results in failed hybridization, are known from a range of plant taxa including maize (Mangelsdorf and Jones, 1926; Kermicle, 2006), gingers (Kay, 2006; Kay and Schemske, 2008; Yost and Kay, 2009), monkeyflowers (Searcy and Macnair, 1990), and tomatoes (Baek *et al.*, 2015). These interactions often act asymmetrically (Tiffin *et al.*, 2001), as we see in this study. Overall, these results indicate that the inability of hybrid formation may be impacted by multiple mechanisms acting in tandem each with variability in symmetry and their overall impact on the likelihood of hybrid formation in nature

A variety of mechanisms, including style length disparity (e.g., Kay, 2006; Baek *et al.*, 2015) and the arrest of pollen tube growth (e.g., Kermicle, 2006), are implicated in these pollen-pistil barriers. Mating system may also play a role; when one hybridizing species is self-incompatible (SI) and the other is self-compatible (SC), SI pollen may pollinate SC ovules, but the reverse cross fails through the production of improperly developed seeds (Brandvain and Haig, 2005; Bedinger *et al.*, 2017). Although both *B. stricta* and *B. retrofracta* are SC, two lines of evidence suggest that this mechanism may play a role. First, average microsatellite-based F_IS_ in *B. stricta* is 0.89 (Song *et al.*, 2006), while average microsatellite-based G_IS_ in *B. retrofracta* is 0.71 (Rushworth *et al.*, 2018). These population statistics are dependent on the level of heterozygosity estimated in each given study, and care should be taken when comparing them. Nonetheless, the large disparity may suggest larger effective population size in *B. retrofracta*, and perhaps higher levels of outcrossing. If this is the case, the SI × SC mechanism predicts that *B. stricta* will succeed only as a maternal parent when paired with *B. retrofracta*, consistent with our results (Figures 2 and 3). Additional support for an SI × SC mechanism would be found in the characterization of aberrant seeds formed by the incompatible cross direction, which is a potential area for further research. Substantial future work is needed to understand the genetic mechanisms underlying hybrid incompatibility in *Boechera*.

Critical to the evolution of hybrid incompatibilities is the environments in which they evolve. Geographic isolation, for example, may strongly impact the evolution of reproductive barriers, leading to expression of barriers in sympatric but not allopatric populations (Coyne and Orr, 2004). This pattern is often seen in the process of reinforcement, where secondary contact leads to increased reproductive isolation between populations (Hopkins, 2013). Although this study is underpowered to detect differences between allopatric and sympatric populations, it is worth noting that all four *B. stricta* lineages that successfully produced hybrids are from populations allopatric with *B. retrofracta*. Additionally, ecological variation will determine patterns of natural selection on hybrids. The suitability of *B. stricta* as a maternal, but not paternal, parent suggests that hybrids are more likely to arise in *B. stricta* habitat. Research focusing on natural selection on *B. stricta* × *B. retrofracta* hybrids should be undertaken in the habitats in which these lineages naturally arise and thrive.

Studies of sexual/asexual dynamics are rarely undertaken in the field, which substantially influences our understanding of the evolution of sex (Neiman *et al.*, 2018). Comparison with studies conducted in the natural environment are thus of vital importance to understanding the real-life costs of sex. Additionally, pairing controlled experiments with those in the field can offer new insight into evolutionary processes (e.g., Anderson *et al.*, 2011). In the field, asexual lineages have higher fitness than sexuals, driven by substantially higher over-winter survival of asexuals, with no evidence for differences in probability of reproduction, fruit number, or fecundity (Rushworth *et al.*, 2019). In contrast, the work presented here showed that de novo hybrids were less likely to reproduce than selfed lineages, resulting in lower overall fitness (Table 1). However, hybrids produced more fruits than sexuals, and had higher variance in fruit number (Figure 4), suggesting that some hybrid lineages may enjoy extremely high fitness. The underlying reasons for this disparity in results are unclear, but if hybrids that form in the wild reproduce sexually and asexually, it is likely that selective patterns between them will differ. For example, if the perceived rarity of wild sexual hybrids is due to their rapid extinction, this may be due to the reproductive incompatibilities reported here. Alternatively, the asexuals in Rushworth *et al.* (2019) may have been recently formed, and thus lacked the accumulated mutations reported in other asexual *Boechera* lineages (Lovell *et al.*, 2017). Further work ascertaining the range of ages for asexual lineages, and the stage at which asexuality occurs, would inform our understanding of selective mechanisms interacting with hybridization and asexuality.

The evolution of sex is strongly influenced by correlated traits that interact uniquely with selective agents in the natural environment. Given the hybrid origins of many asexual lineages, mating system—particularly outcrossing between species—may be particularly important to sexual/asexual dynamics in flowering plants. We found that the expression of multiple reproductively isolating barriers manifest at several life history stages following hybridization, likely impacting the frequency of sexual hybrids in natural populations. Our results contribute to a growing body of literature showing that the origin and fitness of hybrids is interwoven with the costs of sex in the wild.

## Acknowledgements

The authors wish to thank the Mitchell-Olds lab group, especially Baosheng Wang, for access to *B. stricta* chloroplast sequences and germplasm. We also wish to acknowledge Duke University Greenhouses and Phytotron, especially Greg Piotrowski, for their assistance and advice. Dan Runcie and Yaniv Brandvain offered guidance with statistical analyses.

## Data availability

Upon acceptance, primary data on cross success and greenhouse study will be deposited in Dryad. Original sequencing data will be deposited in Genbank and updated in Supplementary Material. Sequencing data from *B. stricta* genotypes is found in Wang *et al.* (2019).

