## Supplementary Material for "The evolution of sex is tempered by costly hybridization in *Boechera* (rock cress)"

**Table S1. Parent lineages used in crosses.** All accessions are of known genotype and were grown for at least one generation in the greenhouse prior to use in crosses. Populations that contain an accession from both *B. stricta* and *B. retrofracta* have both species present and are considered sympatric. Parents that were successful in producing F2s are indicated by an “x” in the column “F2 success.”

| Population | Latitude | Longitude | Accession | Species | F2 success |
|---|---|---|---|---|---|
| MIL | 44.367183 | -113.356783 | MIL7B | *B. retrofracta* | |
| | | | MIL32A | *B. stricta* | |
| TWM | 44.992333 | -113.870033 | TW3 | *B. retrofracta* | |
| | | | TU2-1 | *B. stricta* | |
| YJK | 44.965283 | -114.602050 | YJ7 | *B. retrofracta* | |
| | | | YJ34-1 | *B. stricta* | |
| PKM | 44.604417 | -114.542400 | ES910 | *B. retrofracta* | |
| | | | PK10A.1 | *B. stricta* | |
| RUB | 45.546700 | -113.763200 | RB19A | *B. retrofracta* | |
| | | | RU9B.1 | *B. stricta* | |
| JAM | 44.966883 | -114.085250 | JAM20A | *B. stricta* | x |
| MV9 | 44.182133 | -113.767733 | MV9.4 | *B. stricta* | |
| USL | 44.912217 | -114.386683 | US18A | *B. stricta* | x |
| LTM | 45.705100 | -113.988333 | LTM3#4-1.4 | *B. stricta* | x |
| HES | 39.950700 | -105.593633 | HS12.2-1 | *B. stricta* | x |
| MNK | 44.864833 | -114.297967 | MN8-2 | *B. stricta* | |
| PAN | 45.305567 | -114.382750 | ES913 | *B. retrofracta* | x |
| COR | 45.384033 | -114.704817 | CC10 | *B. retrofracta* | |
| MCK | 44.830300 | -113.943233 | MCK4 | *B. retrofracta* | |
| ALD | 44.807017 | -114.270417 | ALD3 | *B. retrofracta* | |
| SVH | 44.871550 | -114.417550 | SV5 | *B. retrofracta* | |
| BLN | 45.501683 | -114.390600 | BN2 | *B. retrofracta* | |

**Table S2. Accessions used in phylogenetic analysis.** "Taxon" indicates *B. stricta*, *B. retrofracta*, or *B. stricta* × *B. retrofracta* hybrid ("hybrid"). "Source" indicates the original publication for geographic information on accessions, where "Lee" indicates Lee et al. 2011, "Wang" indicates Wang et al. 2019 and "Rushworth" indicates Rushworth et al. 2018. No new sequences were derived for accessions from Wang et al. (2019). Accessions from Rushworth et al. (2018) and Lee et al. (2011) were newly sequenced for t*rn*L.

| Accession | Taxon | Latitude | Longitude | GenBank # | Source |
|---|---|---|---|---|---|
| CR130 | hybrid | 45.556000 | -113.147000 | | Rushworth |
| CR1783 | *B. retrofracta* | 45.009000 | -114.023000 | | Rushworth |
| CR1824 | *B. retrofracta* | 45.118000 | -114.174000 | | Rushworth |
| CR115 | *B. retrofracta* | 44.715000 | -114.348000 | | Rushworth |
| CR136 | *B. retrofracta* | 45.314000 | -114.536000 | | Rushworth |
| CR71 | *B. retrofracta* | 44.965283 | -114.602050 | | Rushworth |
| CR1510 | *B. retrofracta* | 44.871550 | -114.417550 | | Rushworth |
| CR1504 | *B. retrofracta* | 44.367183 | -113.356783 | | Rushworth |
| CR1506 | *B. retrofracta* | 45.384033 | -114.704817 | | Rushworth |
| CR1509 | *B. retrofracta* | 44.830300 | -113.943233 | | Rushworth |
| CR146 | *B. retrofracta* | 45.302000 | -114.338000 | | Rushworth |
| CR1617 | *B. retrofracta* | 45.306000 | -113.889000 | | Rushworth |
| CR197 | *B. retrofracta* | 43.772000 | -114.312000 | | Rushworth |
| CR230 | *B. retrofracta* | 45.787000 | -113.598000 | | Rushworth |
| CR46 | *B. retrofracta* | 44.761000 | -113.196000 | | Rushworth |
| CR686 | *B. retrofracta* | 45.323000 | -113.800000 | | Rushworth |
| CR806 | *B. retrofracta* | 45.576000 | -113.115000 | | Rushworth |
| CR835 | *B. retrofracta* | 44.791000 | -114.253000 | | Rushworth |
| CR87 | *B. retrofracta* | 44.468000 | -113.592000 | | Rushworth |
| CR1315 | *B. retrofracta* | 45.546700 | -113.763200 | | Rushworth |
| CR1331 | *B. retrofracta* | 45.305567 | -114.382750 | | Rushworth |
| CR1321 | *B. retrofracta* | 44.604417 | -114.542400 | | Rushworth |
| CR1548 | *B. retrofracta* | 45.336000 | -113.869000 | | Rushworth |
| CR1538 | *B. retrofracta* | 44.604417 | -114.542400 | | Rushworth |
| CR162 | *B. retrofracta* | 44.974000 | -113.488000 | | Rushworth |
| CR813 | *B. retrofracta* | 45.647000 | -113.069000 | | Rushworth |
| ES913 | *B. retrofracta* | 45.305567 | -114.382750 | | Rushworth |
| CR1524 | *B. stricta* | 39.950700 | -105.593633 | | Lee |
| CR1786 | *B. retrofracta* | 45.443000 | -113.838000 | | Rushworth |
| CR1835 | *B. retrofracta* | 45.386000 | -113.881000 | | Rushworth |
| CR1579 | *B. retrofracta* | 45.007000 | -114.026000 | | Rushworth |
| CR1574 | hybrid | 44.367000 | -113.357000 | | Rushworth |
| CR1273 | *B. retrofracta* | 44.807017 | -114.270417 | | Rushworth |
| CR1516 | *B. stricta* | 44.367183 | -113.356783 | | Lee |
| CR1089 | *B. retrofracta* | 45.501683 | -114.390600 | | Rushworth |
| CR1514 | *B. retrofracta* | 44.992333 | -113.870033 | | Rushworth |
| CR224 | hybrid | 44.406000 | -114.381000 | | Rushworth |
| CR24 | hybrid | 44.948000 | -114.074000 | | Rushworth |
| CR812 | hybrid | 45.489000 | -113.036000 | | Rushworth |

| | | | | | |
|---|---|---|---|---|---|
| CR167 | hybrid | 45.556000 | -113.147000 | | Rushworth |
| CR714 | hybrid | 44.355000 | -113.329000 | | Rushworth |
| CR79 | hybrid | 44.948000 | -114.074000 | | Rushworth |
| MV10 | hybrid | 44.201000 | -113.721000 | | Rushworth |
| CR120 | hybrid | 45.556000 | -113.147000 | | Rushworth |
| CR172 | hybrid | 45.556000 | -113.147000 | | Rushworth |
| CR1165 | *B. stricta* | 44.992333 | -113.870033 | | Lee |
| CR1521 | *B. stricta* | 44.604417 | -114.542400 | | Lee |
| CR826 | hybrid | 44.791000 | -114.253000 | | Rushworth |
| CR118 | hybrid | 45.556000 | -113.147000 | | Rushworth |
| CR144 | hybrid | 44.406000 | -114.381000 | | Rushworth |
| CR56 | hybrid | 44.406000 | -114.381000 | | Rushworth |
| CR60 | hybrid | 44.406000 | -114.381000 | | Rushworth |
| HZ411 | *B. stricta* | 44.794300 | -114.258367 | NA | Wang |
| RP057 | *B. stricta* | 38.692000 | -106.822900 | NA | Wang |
| HZ102 | *B. stricta* | 44.457280 | -114.752880 | NA | Wang |
| RP015 | *B. stricta* | 38.945000 | -106.988400 | NA | Wang |
| RP017 | *B. stricta* | 45.654400 | -113.786700 | NA | Wang |
| RP111 | *B. stricta* | 38.762000 | -106.862800 | NA | Wang |
| RP031 | *B. stricta* | 39.761000 | -110.919400 | NA | Wang |
| RP051 | *B. stricta* | 38.898000 | -107.202600 | NA | Wang |
| HZ053 | *B. stricta* | 45.126580 | -114.136630 | NA | Wang |
| HZ097 | *B. stricta* | 44.443380 | -114.734130 | NA | Wang |
| HZ152 | *B. stricta* | 44.503000 | -114.753200 | NA | Wang |
| RP086 | *B. stricta* | 38.730000 | -106.820100 | NA | Wang |
| RP085 | *B. stricta* | 38.301000 | -107.355800 | NA | Wang |
| RP083 | *B. stricta* | 38.775000 | -106.626900 | NA | Wang |
| RP165 | *B. stricta* | 36.406000 | -112.091700 | NA | Wang |
| RP001 | *B. stricta* | 38.506000 | -109.318900 | NA | Wang |
| HZ449 | *B. stricta* | 44.441583 | -114.733583 | NA | Wang |
| HZ422 | *B. stricta* | 44.832783 | -114.254700 | NA | Wang |
| RP100 | *B. stricta* | 38.510000 | -109.318800 | NA | Wang |
| RP021 | *B. stricta* | 37.913000 | -119.257600 | NA | Wang |
| HZ088 | *B. stricta* | 44.443380 | -114.734130 | NA | Wang |
| CR150 | hybrid | 45.081000 | -114.042000 | | Rushworth |
| CR236 | hybrid | 44.406000 | -114.381000 | | Rushworth |
| CR697 | hybrid | 45.090000 | -114.055000 | | Rushworth |
| CR700 | hybrid | 44.914000 | -114.358000 | | Rushworth |
| CR712 | hybrid | 44.355000 | -113.329000 | | Rushworth |
| CR713 | hybrid | 44.355000 | -113.329000 | | Rushworth |
| CR815 | hybrid | 45.647000 | -113.069000 | | Rushworth |
| CR828 | hybrid | 44.804000 | -114.264000 | | Rushworth |
| CR852 | hybrid | 44.919000 | -114.354000 | | Rushworth |
| CR1525 | *B. stricta* | 44.182133 | -113.767733 | | Lee |
| CR1527 | *B. stricta* | 45.705100 | -113.988333 | | Lee |
| CR1529 | *B. stricta* | 44.912217 | -114.386683 | | Lee |
| RP219 | *B. stricta* | 38.965000 | -112.098800 | NA | Wang |
| CR1520 | *B. stricta* | 44.966883 | -114.085250 | | Lee |
| RP014 | *B. stricta* | 41.844000 | -115.447300 | NA | Wang |

| | | | | | |
|---|---|---|---|---|---|
| RP016 | *B. stricta* | 43.792000 | -114.573700 | NA | Wang |
| RP214 | *B. stricta* | 38.966000 | -111.572200 | NA | Wang |
| RP075 | *B. stricta* | 40.736000 | -110.867600 | NA | Wang |
| RP054 | *B. stricta* | 44.591600 | -114.453300 | NA | Wang |
| RP236 | *B. stricta* | 39.765000 | -110.916300 | NA | Wang |
| HZ015 | *B. stricta* | 44.970670 | -114.085850 | NA | Wang |
| HZ300 | *B. stricta* | 44.792383 | -113.776183 | NA | Wang |
| RP049 | *B. stricta* | 44.941000 | -113.847400 | NA | Wang |
| RP196 | *B. stricta* | 40.823000 | -110.863400 | NA | Wang |
| RP008 | *B. stricta* | 40.435000 | -111.616000 | NA | Wang |
| RP324 | *B. stricta* | 38.211250 | -112.425870 | NA | Wang |
| RP181 | *B. stricta* | 37.612000 | -112.830300 | NA | Wang |
| RP048 | *B. stricta* | 44.616000 | -114.518400 | NA | Wang |
| RP024 | *B. stricta* | 44.355000 | -113.262300 | NA | Wang |
| HZ334 | *B. stricta* | 44.797467 | -113.803217 | NA | Wang |
| HZ337 | *B. stricta* | 44.792600 | -113.778883 | NA | Wang |
| RP012 | *B. stricta* | 40.684000 | -110.931800 | NA | Wang |
| CR133 | hybrid | 44.406000 | -114.381000 | | Rushworth |
| CR62 | hybrid | 44.406000 | -114.381000 | | Rushworth |
| CR715 | hybrid | 44.355000 | -113.329000 | | Rushworth |
| CR77 | hybrid | 44.406000 | -114.381000 | | Rushworth |
| CR97 | hybrid | 45.556000 | -113.147000 | | Rushworth |
| CR99 | hybrid | 44.948000 | -114.074000 | | Rushworth |
| CR1531 | *B. stricta* | 44.864833 | -114.297967 | | Lee |

**Table S3. Hybrids are less likely to reproduce than selfed lines.** Results from GLMM with aborted fruits as the response variable, and plant size as a covariate. Coefficients and standard errors come from model output. Test statistics and *P*-values come from likelihood ratio tests of each term. All *P*-values have been corrected for multiple comparisons. Significant *P*-values are bolded.

| term | condition | df | coef | se | $\chi^2$ | *P* |
|---|---|---|---|---|---|---|
| Cross type | selfed | 1 | 2.80 | 0.659 | 3.29 | 0.21 |
| | hybrid | | 1.54 | 0.417 | | |
| Size | - | 1 | 2.72 | 0.188 | 52.27 | **3.38e-12** |

**Table S4. Hybrids produce more aborted fruits than selfed lines.** Results from GLMM with aborted fruits as the response variable, and plant size as a covariate. Coefficients and standard errors come from model output. Test statistics and *P*-values come from likelihood ratio tests of each term. All *P*-values have been corrected for multiple comparisons. Significant *P*-values are bolded.

| term | condition | df | coef | se | $\chi^2$ | *P* |
|---|---|---|---|---|---|---|
| Cross type | selfed | 1 | 1.21 | 0.202 | 5.09 | 0.096 |
| | hybrid | | 1.83 | 0.209 | | |
| Size | - | 1 | 2.19 | 0.085 | 18.12 | **6.23e-05** |

**Table S5. Fecundity of hybrids and selfed lines does not differ.** Results from GLMM with fecundity as the response variable, and plant size as a covariate. Coefficients and standard errors come from model output. Test statistics and *P*-values come from likelihood ratio tests of each term. All *P*-values have been corrected for multiple comparisons. Significant *P*-values are bolded.

| term | condition | df | coef | se | $\chi^2$ | *P* |
|---|---|---|---|---|---|---|
| Cross type | selfed | 1 | 6.51 | 0.153 | 0.93 | 0.65 |
| | hybrid | | 6.66 | 0.150 | | |
| Size | - | 1 | 6.55 | 0.057 | 0.49 | 0.48 |

**Table S6. Estimated marginal means from all models.** Standard errors are reported on the link-scale from each model.

| *model* | *cross type* | *predicted* | *standard error* | *CI, low* | *CI, high* |
|---|---|---|---|---|---|
| reproduction | hybrid | 0.84 | 0.42 | 0.69 | 0.92 |
| | selfed | 0.95 | 0.51 | 0.87 | 0.98 |
| fruit number | hybrid | 22.41 | 0.17 | 16.17 | 31.05 |
| | selfed | 8.60 | 0.16 | 6.34 | 11.68 |
| fecundity | hybrid | 679.58 | 0.16 | 500.65 | 922.44 |
| | selfed | 787.78 | 0.15 | 591.07 | 1049.95 |
| aborted fruit number | hybrid | 6.21 | 0.2 | 4.18 | 9.23 |
| | selfed | 3.35 | 0.2 | 2.27 | 4.96 |

**Table S7.** Variance estimates and standard deviations for random effects from all models.

| *Model* | *Group* | *Variance* | *SD* |
|---|---|---|---|
| reproduction | crosstype:parent | 0.445 | 0.667 |
| | block | 0.034 | 0.185 |
| fruit number | parent | 1.37e-08 | 1.17e-04 |
| | crosstype:parent | 6.38e-02 | 0.253 |
| | block | 1.14e-02 | 0.107 |
| aborted fruit number | parent | 3.17e-02 | 0.178 |
| | crosstype:parent | 1.15e-02 | 0.107 |
| | genotype | 3.42e-02 | 0.185 |
| | block | 4.92e-02 | 0.222 |
| fecundity | parent | 2.55e-02 | 0.160 |
| | crosstype:parent | 2.94e-02 | 0.172 |
| | block | 2.26e-02 | 0.150 |
| total fitness | parent | 4.39e-03 | 0.066 |
| | crosstype:parent | 5.07e-05 | 0.007 |
| | genotype | 2.14e-02 | 0.146 |
| | block | 0.142 | 0.377 |

**Figure S1. Fecundity does not differ between hybrid and selfed lineages.**
Estimated marginal means from GLMM. Bars show 95% confidence intervals.

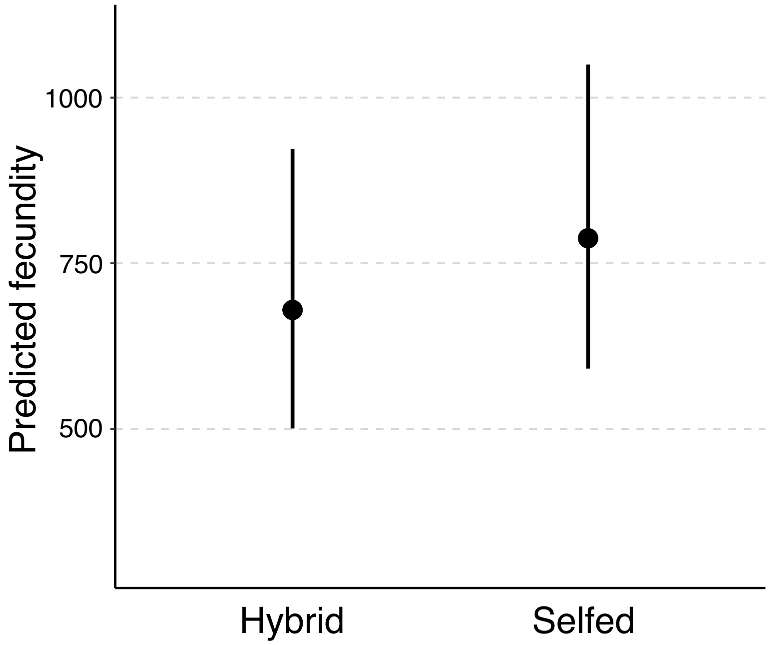

**Figure S2. Selfed lineages are larger than hybrids.** Each data point is raw data for an experimental individual.

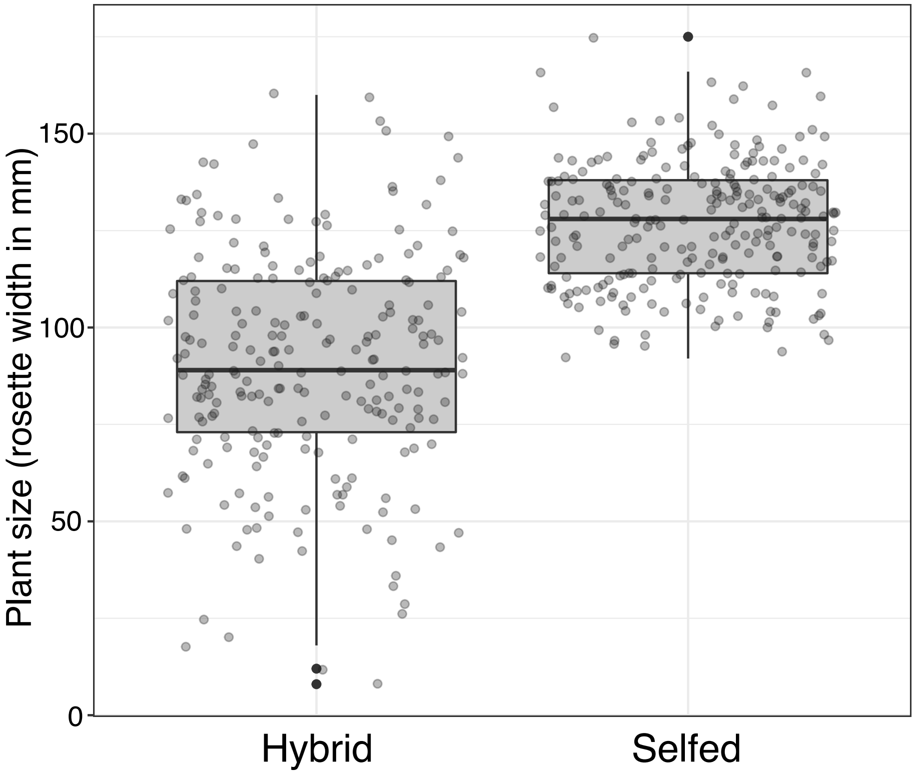
